# An improved MS2-MCP imaging system with minimal perturbation of mRNA stability

**DOI:** 10.1101/2022.02.05.479257

**Authors:** Weihan Li, Anna Maekiniemi, Hanae Sato, Christof Osman, Robert H. Singer

**Author notes:** Correspondence: Robert H. Singer. These authors contribute equally to this work (Weihan Li, Anna Maekiniemi).

## Abstract

The MS2-MCP imaging system is widely used to study the mRNA spatial distribution in living cells. Here, we report that the MS2-MCP system may destabilize the tagged mRNA by targeting it to the nonsense-mediated mRNA decay pathway. We introduce an improved version, which has minimal perturbation of the mRNA stability.

## Main

The spatial distribution of mRNA regulates its gene expression^1^. The MS2-MCP system is widely used to image mRNAs with high spatial-temporal resolution in living cells^1,2^. Specifically, an mRNA of interest is genetically fused to an array of MS2 binding sites (MBS) in its 3’ UTR. Fluorescently labeled MS2-coat proteins (MCP) bind to the MBS array, making the mRNA molecules appear as diffraction-limited spots under wide-field epifluorescence microscopy^3^. Capturing the mRNAs’ innate behavior requires minimum perturbation from the imaging system. Recent advances in the design of the MBS array improved its imaging accuracy by avoiding the aggregation of decay-resistant fragments^3^. In this study, we report that the MS2-MCP system may destabilize the tagged transcript through nonsense-mediated mRNA decay (NMD). We present an improved MS2-MCP system that minimizes the mRNA destabilization.

The length of 3’ UTR regulates mRNA stability. Long 3’ UTRs cause aberrant translation termination and render the mRNA degraded through NMD^4,5^, which is a quality control pathway that degrades mRNAs with premature stop codons, long 3’ UTRs, or upstream open reading frames (uORFs)^6^. The median length of yeast 3’ UTRs is 104 nt^7^, whereas a 24xMBS array is 1,660 nt. Thus, MBS tagging in yeast extends 3’ UTRs by 16-fold on average and, as a result, could render the mRNA a target of NMD. To explore this notion, we genetically fused the MBS array to the 3’ UTRs of eight yeast genes and examined their steady-state mRNA levels using qPCR. Three of the genes, *ATP2, HAC1*, and *PMA1*, had different degrees of mRNA reduction when tagged with the MBS array (Figure 1A). In contrast, the mRNA levels of the remaining five genes, including *ACT1*, were not significantly affected by the MBS tagging (Figure 1A, Supplementary Figure 1A-C) (see Discussion). *ATP2* encodes the ß subunit of F^1^-ATP synthase^8^. Using qPCR and single-molecule RNA fluorescent *in situ* hybridization (smFISH), we found that the level of the *ATP2* mRNA reduced to 30% after MBS tagging, representing the largest mRNA reduction among the genes we tested (Figure 1B-D). The effect of mRNA destabilization did not change when the MBS array was inserted at a different location in the 3’ UTR, the number of MBS repeats was reduced from 24 to 12, or the binding protein MCP-GFP was co-expressed (Figure 1B, Supplementary Figure 2). The destabilization of *ATP2* mRNA exhibited a phenotypic impact as the yeast cells with the MBS-tagged *ATP2* grew slower on a non-fermentable carbon source (Figure 1E). This is consistent with *ATP2*’s role in oxidative phosphorylation. *HAC1* encodes the transcription factor regulating protein folding homeostasis in the endoplasmic reticulum (ER)^9^. The level of the *HAC1* mRNA reduced to 50% after MBS tagging, thereby decreasing the cellular fitness under ER protein folding stress conditions (Figure 1A and F). To test whether the NMD pathway was responsible for destabilizing the MBS-tagged mRNAs, we deleted *UPF1*, a core mediator of NMD^10^. Deletion of *UPF1* recovered the levels of the tagged mRNAs (Figure 1A) and the growth phenotypes (Figure 1E and F). Hence, tagging with an MBS array may destabilize yeast mRNAs through NMD.

**Figure 1.**
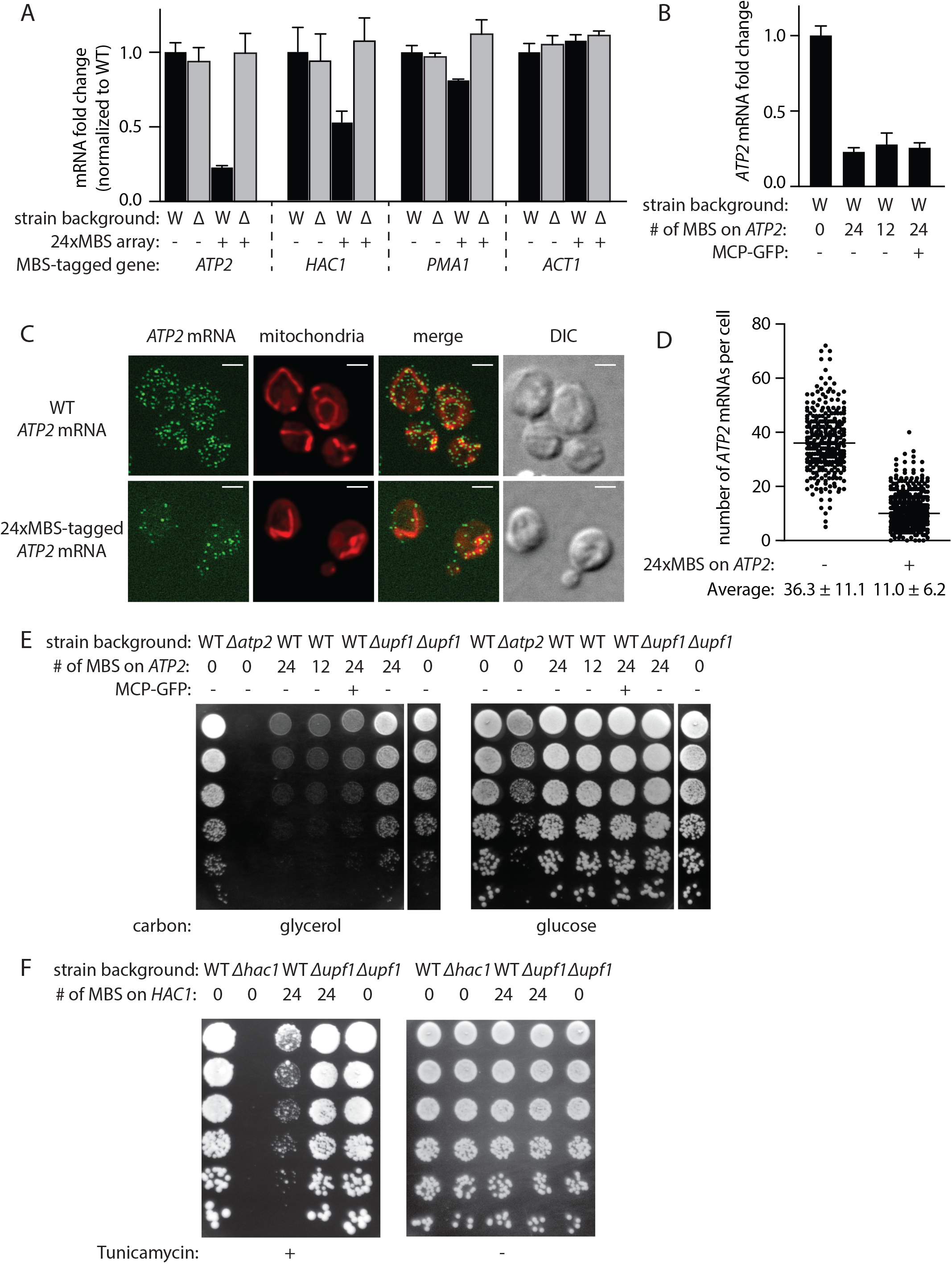
MBS tagging may destabilize mRNA through the NMD pathway in yeast. (A) qPCR of the MBS-tagged mRNAs in WT (W) or Δ*upf1* (Δ) strain backgrounds. The mRNA levels were normalized to their corresponding WT mRNAs. (B) qPCR of the *ATP2* mRNAs tagged with 24x or 12x MBS array in the presence or absence of MCP-GFP. (C) Representative images of *ATP2* mRNA smFISH (green). Mitochondria were labeled using mitochondrial-targeted fluorescent protein (red). Images were max-Z projected. Scale bars are 2 µm. (D) The number of *ATP2* mRNAs per cell as quantified from the smFISH images. (E, F) Cell growth assay on plates with the indicated carbon source (E) or the ER stress inducer tunicamycin (F).

Next, we sought to modify the MS2-MCP system to minimize the destabilization effect. Previous studies showed that long 3’ UTRs in yeast cause inefficient translation termination and subsequently trigger NMD^4^. Tethering the yeast translation termination factor, SUP35, or Poly(A) binding protein, PAB1, to the vicinity of the stop codon increased the efficiency of translation termination, thereby protecting the mRNA from NMD^4,11^. Inspired by these studies, we expressed the fusion protein MCP-GFP-SUP35 or MCP-GFP-PAB1, which should bind to the MBS-tagged mRNA, making the mRNA fluorescent through GFP and correcting the mRNA stability through SUP35 or PAB1 (Figure 2A). When using MCP-GFP-PAB1, we noticed that the MBS-tagged mRNAs were not detected using fluorescent microscopy (Supplementary Figure 3A). We surmised that the MCP-GFP-PAB1 bound to the poly(A) tails of other mRNAs through the PAB1 RNA-binding domain. To test this idea, we used PAB1(F170V, F366V) (referred to as PAB1* in this study), which is a mutant that harbors two point mutations at conserved residues of its RNA-binding domain (Supplementary Figure 3B, C). The RNA binding affinity of PAB1* is nearly 1,000-fold lower than that of the wild-type (WT)^12^. As a result, the MCP-GFP-PAB1* is expected to have higher specificity for the MBS arrays. Indeed, MBS-tagged mRNAs became visible when using MCP-GFP-PAB1* (Supplementary Figure 3A). Expressing the fusion protein MCP-GFP-SUP35 or MCP-GFP-PAB1* recovered the steady-state levels of the MBS-tagged *ATP2, HAC1*, and *PMA1* mRNAs, as well as their corresponding growth phenotypes (Figure 2B-D). *ACT1* represents the mRNAs that were not destabilized by the MBS tagging. The MCP fusion proteins didn’t change the level of the tagged *ACT1* mRNA (Supplementary Figure 4). Therefore, these data showed that the MCP-GFP-SUP35 and MCP-GFP-PAB1* can be used to image mRNAs with minimal perturbation of mRNA stability.

**Figure 2.**
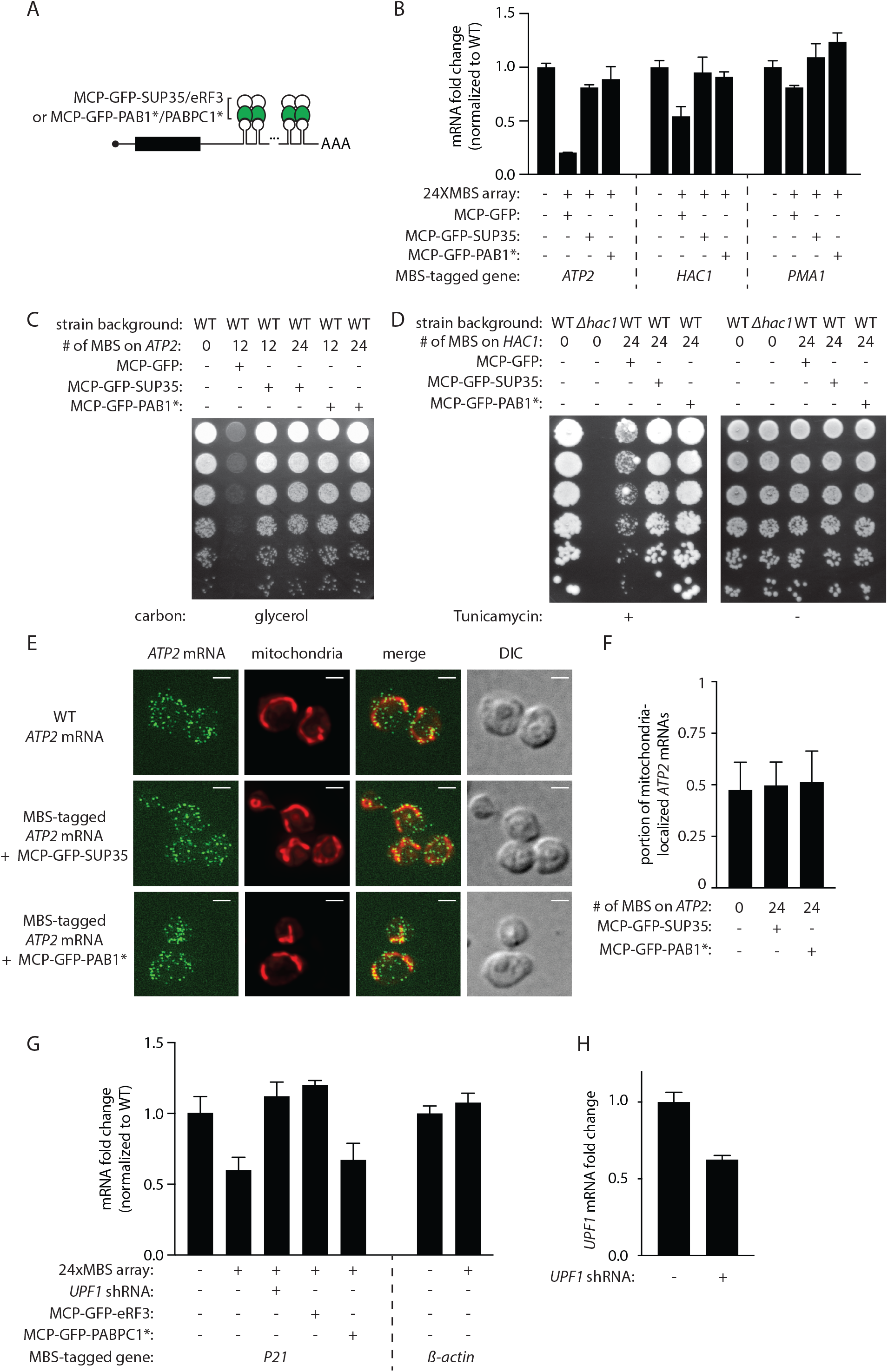
The improved MS2-MCP system minimizes the mRNA destabilization in yeast and mammalian cells. (A) Cartoon illustration of the improved MS2-MCP system. (B) qPCR of the indicated mRNAs. The mRNA levels were normalized to their corresponding WT mRNAs. (C, D) Cell growth assay with the same conditions as in Figure 1E and F. (E) smFISH of *ATP2* mRNAs. The experimental setups and scale bars are the same as in Figure 1C. (F) Quantifying the proportion of the mitochondria-localized *ATP2* mRNAs from the smFISH images. (G) qPCR of the MBS-tagged *P21* mRNA in U2OS cells and *ß-actin* mRNA in MEFs. The mRNA levels were normalized to their corresponding WT mRNAs. Error bars were calculated from qPCR replicates. (H) qPCR of *UPF1* mRNA in the presence or absence of the *UPF1* shRNA.

We tested whether the improved MS2-MCP system could faithfully report on mRNA distribution. Previous studies showed that the *ATP2* mRNA is translated at mitochondrial periphery, thereby promoting the import of the nascent ATP2 proteins^13^. In fixed cells, we imaged *ATP2* mRNAs using smFISH, and mitochondria using a mitochondrial-targeted fluorescent protein. About half of the *ATP2* mRNAs co-localized with mitochondria in WT cells, as well as in cells expressing the improved MS2-MCP system (Figure 2E and F). This result suggested that the improved MS2-MCP system did not perturb the mRNA localization. In living cells, we imaged *ATP2* mRNAs using the MCP-GFP-SUP35 and showed that *ATP2* mRNAs moved in trajectories that coincide with the mitochondrial network (Supplementary Figure 5, Supplementary Movie 1 and 2). Therefore, the improved MS2-MCP system faithfully reported on the mRNA distribution.

To extend our study to mammalian cells, we examined the *ß-actin* mRNA in mouse embryonic fibroblasts (MEFs) and *P21* mRNAs in U2OS cell lines because they have been tagged with the MBS arrays in previous studies^14,15^. MBS tagging reduced the *P21* mRNA level but did not change the *ß-actin* mRNA level (Figure 2G). This is similar to what we observed in yeast as the MBS-induced destabilization varied from gene to gene. When *UPF1* was knocked down using shRNA, the level of the tagged *P21* mRNA recovered, suggesting that the mRNA destabilization occurred through the NMD pathway (Figure 2G, H). To minimize the destabilization, MCP-GFP was genetically fused to eRF3 or PABPC1(F142V, F337V) (referred to as PABPC1* in this study), which are the mammalian homologs of SUP35 and PAB1*. Expressing the MCP-GFP-eRF3 fully restored the level of the *P21* mRNA, showing the fusion protein’s conserved function in yeast and mammalian cells. On the other hand, expressing the MCP-GFP-PABPC1* moderately restored the mRNA level (Figure 2G). The degree of mRNA recovery increased as the expression level of the MCP-GFP-PABPC1* increased (Supplementary Figure 6). Therefore, MCP-GFP-eRF3 could better restore the stability of the MBS-tagged *P21* mRNA in mammalian cells.

In summary, we reported that the MBS tagging may decrease the mRNA stability by targeting the transcripts to the NMD pathway. We presented an improved MS2-MCP system, in which MCP-GFP-SUP35/eRF3 or MCP-GFP-PAB1*/PABPC1* protected the tagged yeast/mammalian mRNAs from NMD. Orthogonal RNA imaging methods, including the PP7-PCP system or aptamer-based RNA imaging methods, achieve single-molecule resolution using a similar strategy of repeated tags^16^. These tags increase the length of the 3’ UTR and may analogously destabilize the mRNA. Our improved MS2-MCP system provides a template to antagonize NMD and correct the mRNA stability. In this study, we observed that the MBS-induced NMD varied from gene to gene even though the tagged mRNAs had similar 3’ UTR lengths. This resembles the situation in mammalian cells where NMD may differentially regulate endogenous mRNAs with comparable 3’ UTR lengths. The differential regulation by NMD may be determined by additional factors, including cis-elements in the 3’ UTR, proteins bound to the mRNA, and the translation readthrough rate^17-21^. We recommend to assess the effect of MBS tagging on an mRNA of interest before choosing the MS2-coat protein construct. If MBS tagging does not change the mRNA stability, one can use the original MCP-GFP. In contrast, if MBS tagging reduces the mRNA stability, we recommend the MCP-GFP-SUP35/eRF3 or MCP-GFP-PAB1*/PABPC1*. Altogether, the new MS2-MCP system has minimized impact on mRNA stability and can faithfully report on the mRNA’s spatial distributions in living cells. We believe it will see widespread applications in future studies.

## Acknowledgements

We thank Allan Jacobson, U. Thomas Meier, Robert A. Coleman, and members of the Singer lab for their insightful discussions. This work was supported by American Heart Association Postdoctoral Fellowship #903024 (WL), 1R35 GM136296-01 (RHS), and the European Research Council ERCStG-714739 IlluMitoDNA (CO).

---

**Supplementary Figure 1.**
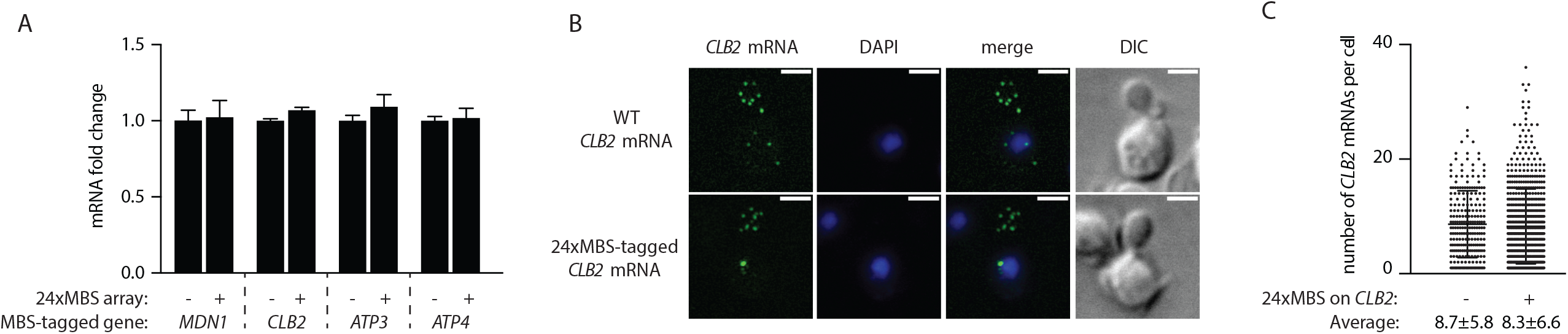
MBS tagging does not destabilize all mRNAs. A) qPCR of untagged and MBS-tagged mRNAs. The mRNA levels were normalized to their corresponding WT mRNAs. (B) Representative images of *CLB2* mRNA smFISH (green). DNA was stained with DAPI (blue). Scale bars are 2 µm. (C) The number of *CLB2* mRNAs per cell quantified from smFISH images.

**Supplementary Figure 2.**
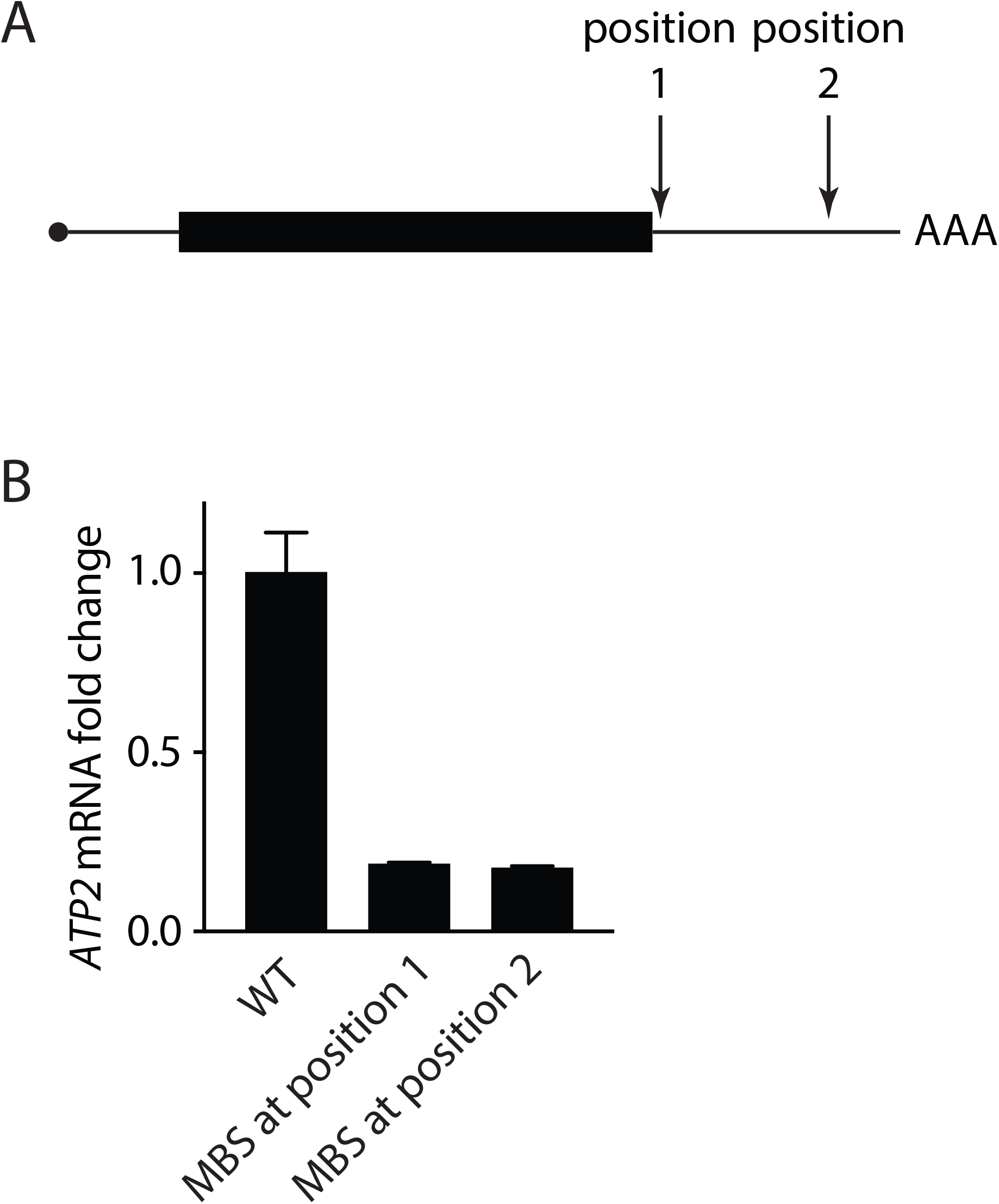
The steady-state levels of the *ATP2* mRNA remained the same after altering the location where the MBS array was inserted. (A) Illustration of the tagging positions. Position 1 is immediately downstream of the stop codon, while position 2 is 162 nt downstream of the stop codon. (B) qPCR of the indicated *ATP2* mRNAs.

**Supplementary Figure 3.**
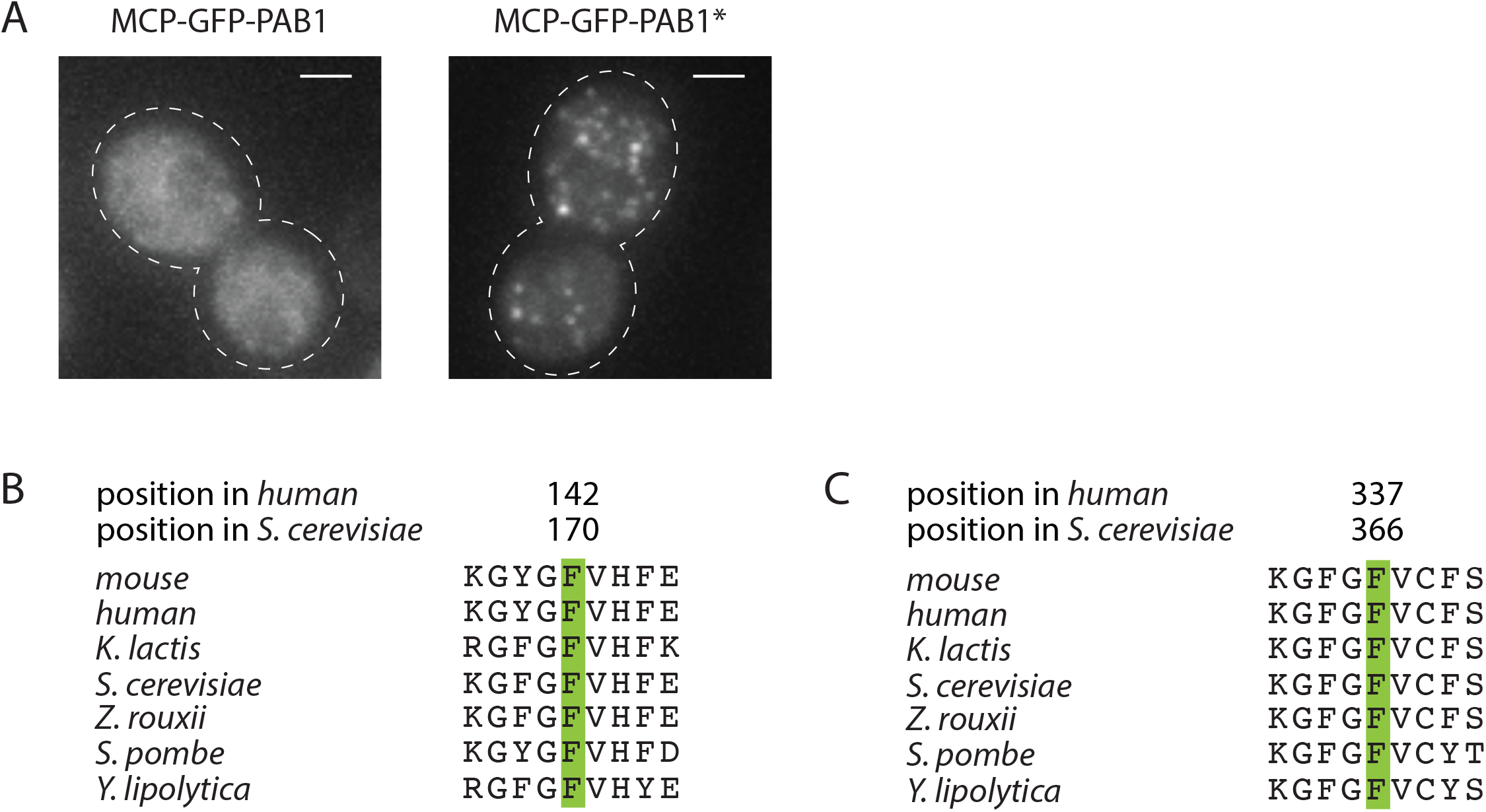
MCP-GFP-PAB1* can be used to image the MBS-tagged *ATP2* mRNAs. (A) Live-cell imaging of yeast cells expressing MCP-GFP-PAB1 (left) or MCP-GFP-PAB1* (right). Images were max-Z projected. Scale bars are 2 µm. Cell outline is marked with white dashed line. (B, C) Sequence alignment of the PAB1 homologs flanking the two conserved phenylalanines, which are the F170 and F366 in the yeast PAB1 (F142 and F337 in the human PABPC1). These two phenylalanines were mutated to valines in PAB1*/PABPC1*.

**Supplementary Figure 4.**
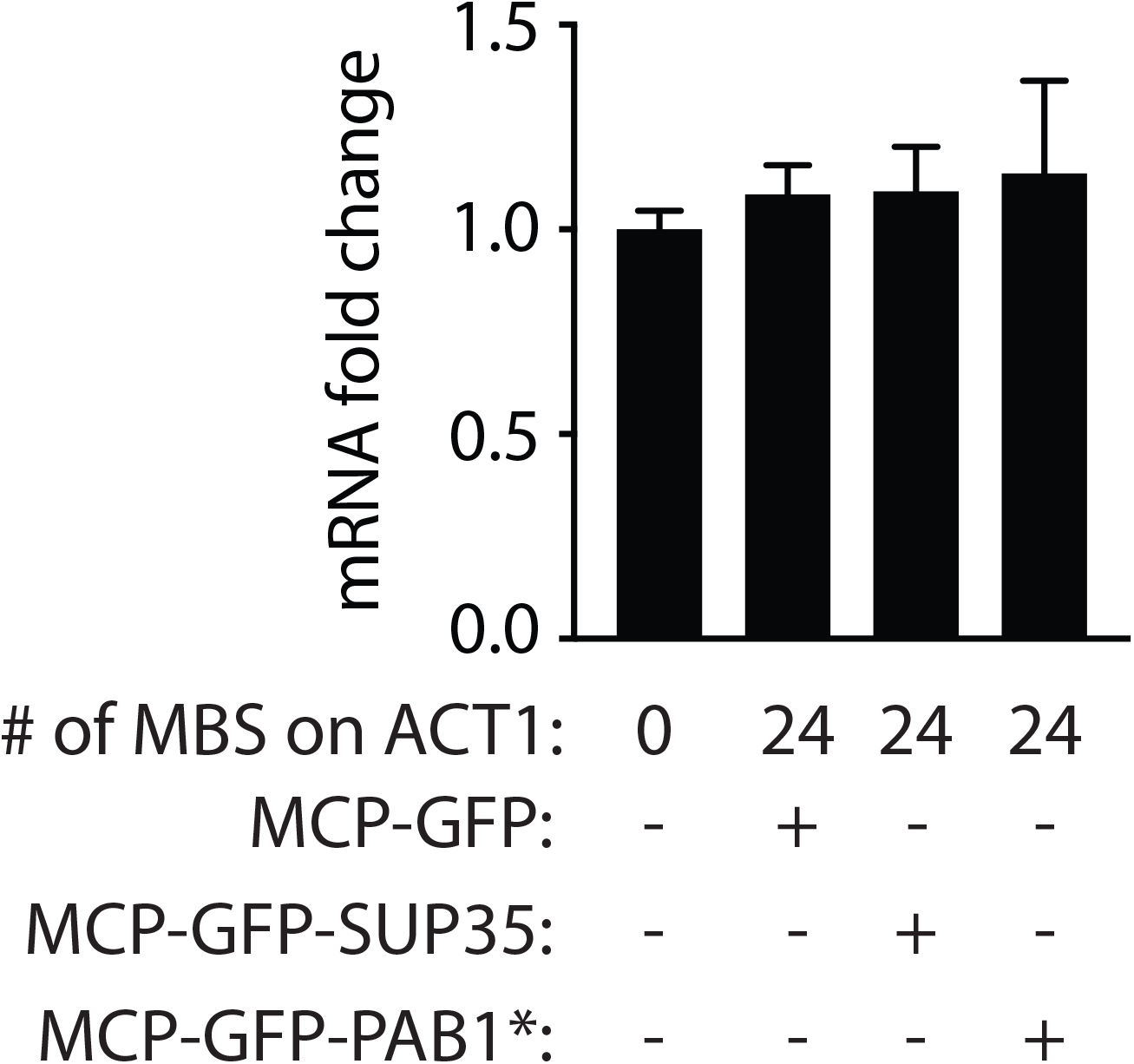
Expressing MCP-GFP-SUP35 or MCP-GFP-PAB1* did not change the steady-state level of the MBS-tagged *ACT1* mRNA. The indicated *ACT1* mRNAs were measured by qPCR.

**Supplementary Figure 5.**
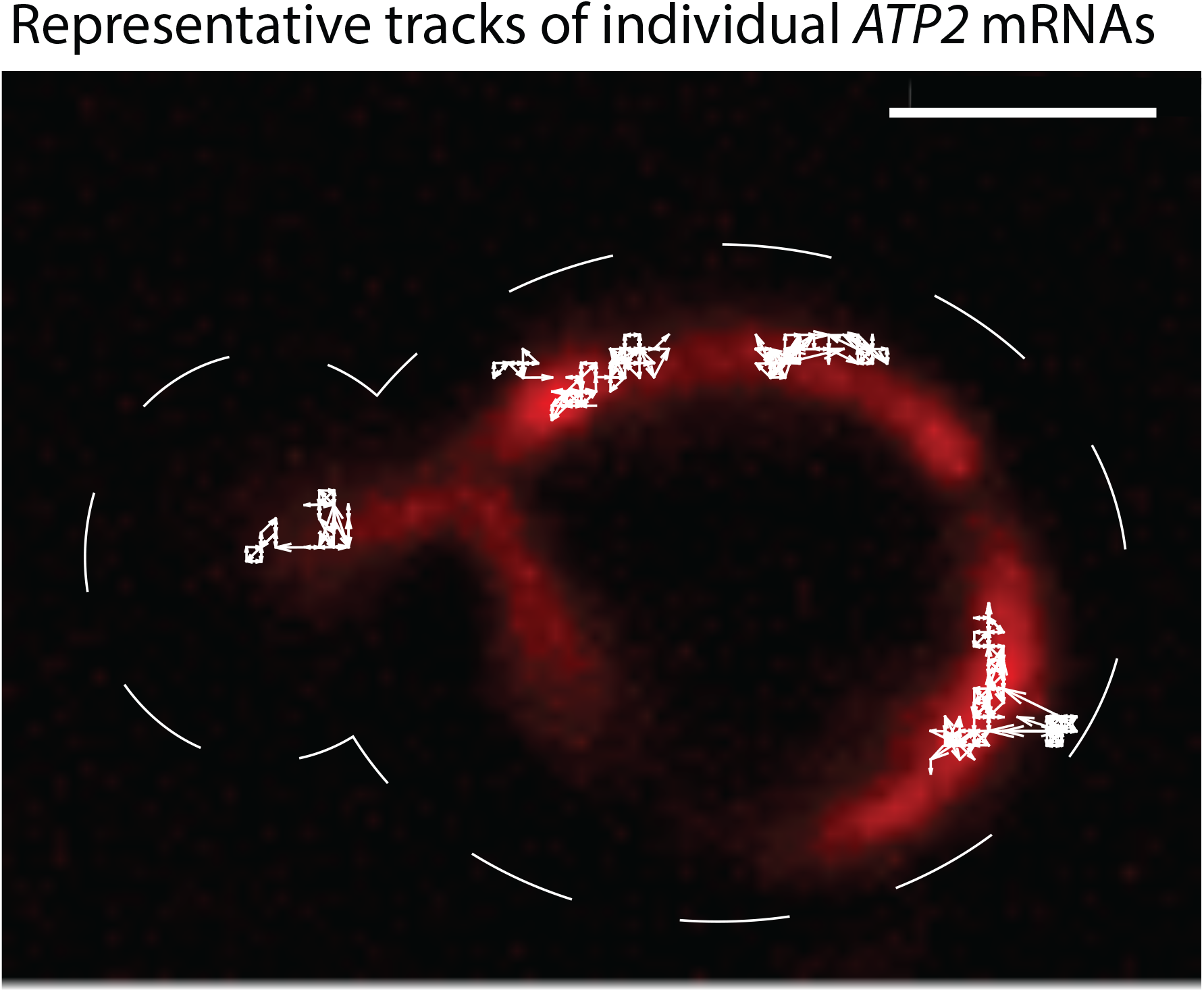
Tracking single-molecule *ATP2* mRNAs using the improved MS2-MCP system. The original movie is shown in Supplementary Movie 1. Mitochondria were labeled using a mitochondria-targeted mKate2. MBS-tagged *ATP2* mRNAs were imaged with MCP-GFP-SUP35. The movie was acquired at 20 frames per second in one z plane. The figure shows the molecular trajectories of the mRNAs that were tracked for at least 40 consecutive frames (white arrowed lines). Mitochondria are in red. Cell outline is marked with white dashed line. Scale bar is 2 µm

**Supplementary Figure 6.**
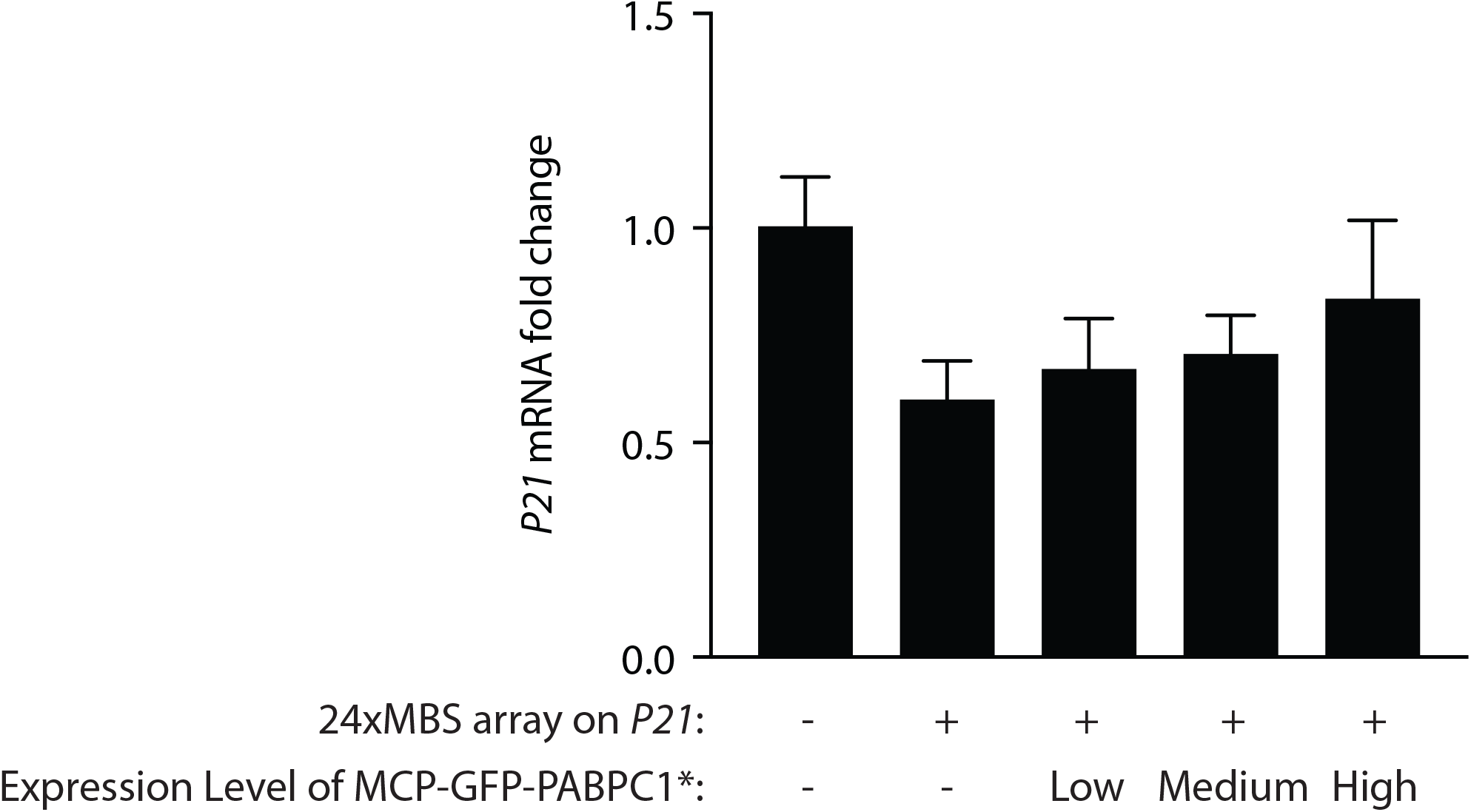
The *P21* mRNA recovery increased as the expression level of the MCP-GFP-PABPC1* increased. Cells were sorted by FACS based on the expression level of MCP-GFP-PABPC1*. The level of the *P21* mRNA was examined by qPCR. Error bars were calculated from qPCR replicates.

**Supplementary Movie 1. Live-cell imaging of *ATP2* mRNAs using MCP-GFP-SUP35**. Mitochondria (red) were labeled using a mitochondria-targeted mKate2. MBS-tagged *ATP2* mRNAs (green) were imaged with MCP-GFP-SUP35. The movie was acquired at 20 frames per second in one z plane. Scale bar is 2 µm.

**Supplementary Movie 2. Live-cell imaging of *ATP2* mRNAs using MCP-GFP-SUP35**. Experimental setup is the same as Supplementary Movie 1. Scale bar is 2 µm.

